# Root expansion induced by Sulfur limitation and mild heatwaves mitigated yield loss under a severe heatwave in grasslands

**DOI:** 10.64898/2025.12.02.691764

**Authors:** Andreu Cera, Servane Lemauviel-Lavenant, Quentin Dupas, Sophie Brunel-Muguet

## Abstract

Grasslands, like many temperate ecosystems, are threatened by heatwaves which have been shown to be more frequent. Plant diversity can buffer yield loss due to heatwaves, but this positive effect may be modified by nutrient availability. Soil sulfur (S) has declined over the last decades and although it is crucial for coping with abiotic stresses, its role in grassland responses to heatwaves remains poorly documented. We aimed to determine how S nutrition modulates the responses of grasslands to heatwaves. Four monocultures and two mixtures were grown under two S levels and exposed to four thermoprotocols: i) control, ii) two successive mild heatwaves, iii) one severe heatwave, and iv) a recurrent sequence combining mild and severe heatwaves. We measured biomasses, leaf S compounds, δ^13^C as a proxy of water-use efficiency, leaf temperature, and photosystem II efficiency. S nutrition interacted with thermoprotocols. Under a severe heatwave, standard S promoted higher shoot production and better water-use efficiency. In contrast, under recurrent events, S limitation enhanced root:shoot ratio, water-use efficiency, shoot growth, and led to lower leaf temperature. S-containing metabolites did not vary with heatwaves, suggesting no direct metabolic. Under recurrent heatwaves, S limitation could indirectly improve tolerance to severe heatwaves.

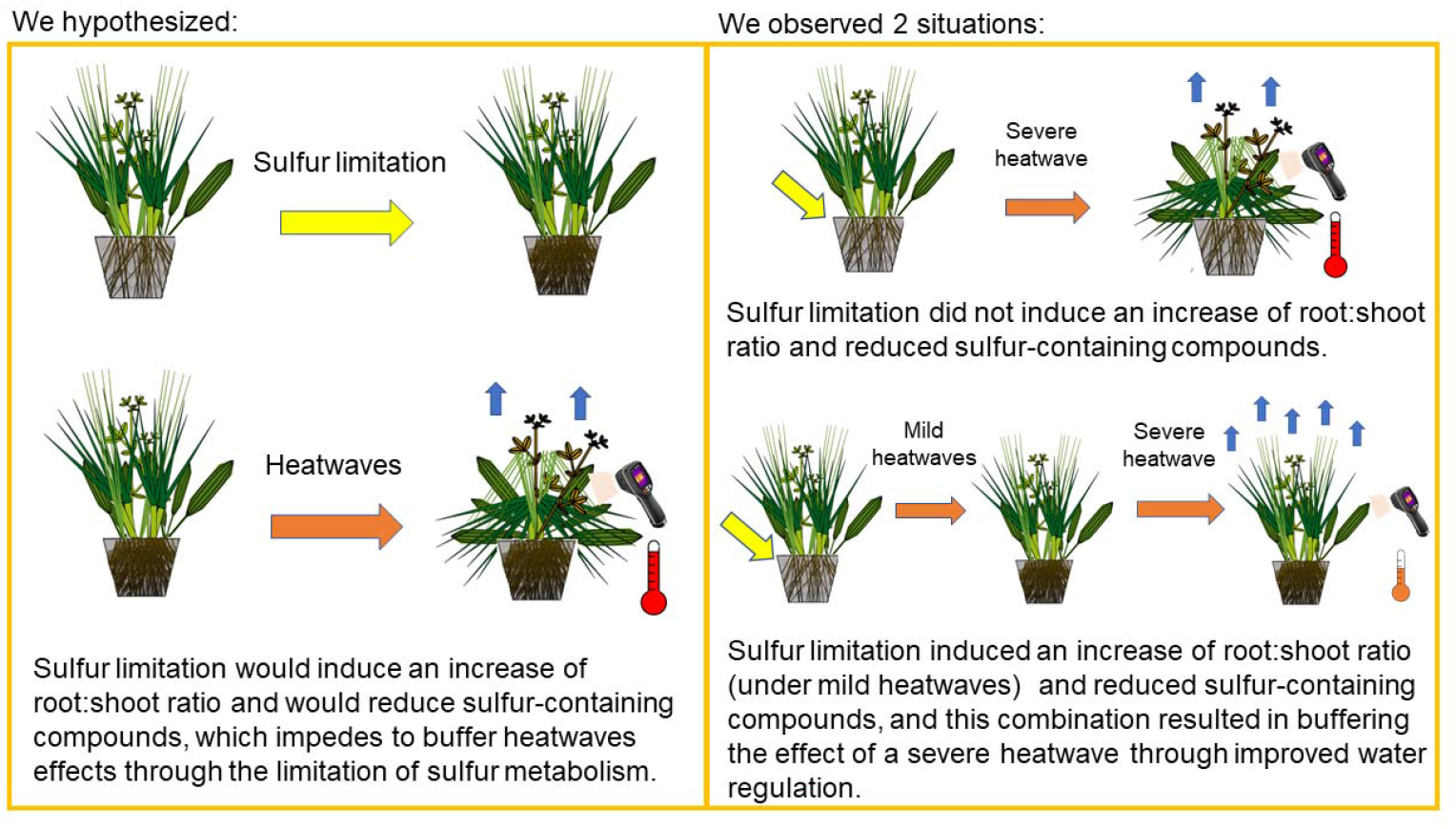

## Introduction

Grasslands are the primary land use in Europe’s agricultural area; they provide many ecosystem services, from fodder supply to biodiversity conservation. Over decades, land-use change and the intensification of management have led to the loss of high diversity grassland areas and a decrease in their multifunctionality (Schils *et al*., 2022). Compounding these pressures, global change threatens the functioning of grasslands and their stability. Among the major threats, heatwaves are expected to be more frequent, longer and more intense in most parts of the world (IPCC reports), posing increasing risks to temperate grassland productivity (Wang *et al*., 2008; Langworthy *et al*., 2020). Grasslands represent a solution for sustainable livestock production (Dumont *et al*., 2019). Adapting them to global changes represents a major challenge in addressing the livestock crisis (Bilotto *et al*., 2024).

Grassland species have developed mechanisms to cope with heatwaves. These mechanisms are linked to the avoidance of excess energy (Körner, 2013), as exemplified by leaf cooling (Urban *et al*., 2017), or the reduction of heat stress by osmoprotectants or antioxidant metabolites (Wahid *et al*., 2007). The timing of heatwaves plays a crucial role in determining their impact (Jagadish *et al*., 2021). The season of occurrence determines the intensity and the effects of heatwaves: mild heatwaves, more likely to occur in spring, have a relatively neutral effect on production in grassland species (De Boeck *et al*., 2011), whereas more drastic summer heatwaves cause more severe damage (Wang *et al*., 2008). Most studies have focused on how intensity and duration of a single and isolated high temperature event modulate plant performances. Nevertheless, heatwaves are repeated and erratic events rather than sustained stressful periods (Magno Massuia de Almeida *et al*., 2021), indicating that their frequency must be considered to realistically characterize their effects on plant growth and functioning. Plants have the ability to acclimate to these recurrent events via memory-related processes (Ogle *et al*., 2015). They can store and retrieve information they have acquired from previous exposures, and acclimate to the next event as through phenotypic plasticity (Gessler *et al*., 2020). In addition to these individual effects, growth in a plant community imply plant-plant interactions that can exert a key role in response to climatic disturbances. Plant communities, constituted by different species that do not share the same resource use strategy nor resource requirements, favor mechanisms such as niche complementarity or facilitation, which enable plant mixture a better response to heatwaves than monocultures (Grant *et al*., 2014). Cera et al. (2025a) demonstrated for grasslands that mixtures were more productive than the average of monocultures, experiencing overyielding under single and recurrent heatwaves. Hence, the study of the effect of plant diversity under climatic disturbances or other abiotic factors, as well as the analysis of the mechanisms involved, are relevant for adapting grasslands to future challenges linked to climate change.

Sulfur (S) nutrition in grasslands has been debated for a long time in Europe due to growing limitations in soil since the 1980s (Scherer, 2001; Seaton *et al*., 2023). S is an essential macronutrient for growth but plays also a key role in plant responses to changes in abiotic factors (Hawkesford *et al*., 2012). Mass production is reduced under S limitation for many grassland species, but may be accompanied by an increase in their root:shoot mass ratio (Gilbert and Robson, 1984). In response to abiotic factors, S metabolism of plants is modified; for example, they increase the synthesis of glutathione or other S-containing compounds derived from sulphate metabolism (Kopriva and Rennenberg, 2004; Hasanuzzaman *et al*., 2018). From the observed outcomes at the individual level, S limitation can result to changes in the plant community composition, as it modifies the species productivity and competitive abilities. Tallec et al. (2008) compared two grassland species, *Lolium perenne* and *Trifolium repens*, on the basis of their S requirements. They observed that *T. repens* performed worse under S-limiting conditions, while *L. perenne* was unaffected. In Fabaceae, sulfur deficiency limit N_2_ fixation and thus performances through both S and S metabolisms (Varin et al. 2010). Consequently, S-limiting conditions lead to lower proportions of legumes in the community. However, little is known about how S availability influences plant community responses under heatwaves. To address this knowledge gap, our study investigated how S nutrition modulates the responses of grassland monocultures and mixtures to single and recurrent heatwaves. Specifically, we grew four monocultures and two mixtures under two contrasting S levels (standard: 0.5 mM; low: 0.1 mM) and subjected them to four thermoprotocols (control, mild heatwaves, severe heatwave, recurrent heatwaves). We measured shoot and root masses, leaf contents of Sulfur and S-related compounds (sulfate and glutathione), δ^1 3^C as a proxy of water-use efficiency, and heat-sensitive parameters such as leaf temperature and photosystem II efficiency. We hypothesized that: (i) S limitation would reduce shoot mass and increase root:shoot mass ratio; (ii) S limitation would constrain S metabolism, reducing concentrations of S-containing compounds, limiting protective mechanisms against heatwaves; (iii) as a consequence of (ii) plants under low S would show higher leaf temperature, lower water-use efficiency, and reduced PSII efficiency; and (iv) these effects would be more pronounced in mixtures under low S due to species-specific differences on Sulfur requirements.

## Material and methods

### Species and varieties used

This experiment was conducted in a greenhouse at Université de Caen-Normandie, France (49°11′09 N, 0°21′32 W), using temperate grassland species commonly sown in Europe (Cera *et al*., 2025, Preprint): *Lolium perenne* L., *Festuca rubra* L., *Lotus corniculatus* L., and *Plantago lanceolata* L. The varieties employed were “Elixir” for *L. perenne* (provided by Cerience), and “Bardance” for *F. rubra*, “Lotar” for *L. corniculatus*, and “Captain” for *P. lanceolata* (all supplied by Barenbrug).

### Culture preparation and management

Seeds of each species were germinated separately on November 24^th^, 2023, in germination trays with a mix of osmotic water and perlite (3:1, v:v), under greenhouse conditions at 20°C/16°C (day/night) with a photoperiod of 16 hours of day under artificial lightening (sodium high pressure ‘phytoclaude 400 W’). On December 4th, 2023, seedlings were transplanted into 4 L cylindrical pots, filled with quarry silica sand (Silica BB 0.8-1.8 DS:BL, Sibelco, Mios, France). We prepared three types of grassland mixtures to study the effect of species diversity on grassland performances: monocultures for each of the four species; a grass-legume mixture of *Lolium perenne* and *Lotus corniculatus*; and a mixture of the four species. In each pot, 2 to 3 seedlings were placed into each of the 8 pre-marked holes to establish a total of 8 plants per pot. Fifteen days after transplantation, thinning was performed to keep only one individual per hole. We applied three mowing at 5 cm to all pots at three dates: March 11^th^, 2024; May 6^th^, 2024; and the last mowing and harvest on July 1^st^, 2024 to simulate mowing in the local context (Normandy, France). Pots were placed on a mobile rack system and repositioned monthly to minimize microhabitat effects. Pests, mainly aphids and thrips, were controlled by spraying or washing the leaves with tap water.

### Thermoprotocols

To study the effects of heatwaves, we designed four distinct thermoprotocols, as tested in (Cera *et al*., 2025, Preprint). From germination until final harvest, all plants were grown in a greenhouse under a standard thermoperiod of 20/16°C (16h day/ 8h night). Sixteen weeks after transplanting the plants into sand pots, on March 11^th^, 2024, we assigned the pots to one of the four thermoprotocols: (i) Control, where plants remained under the standard thermoperiod; (ii) Mild heatwaves, involving two periods of moderately elevated temperatures (25/18°C, day/night), each lasting five days and occurring 16 and 18 weeks after transplantation; (iii) Severe heatwave, consisting of a single five-day event at high temperatures (41.5/19.5°C, day/night) starting 26 weeks after transplantation; and (iv) Recurrent heatwaves, where plants experienced the three heatwave events (two mild and one severe). Heatwave events were applied to the plants in a separate greenhouse compartment. The temperature regimes in both control and heatwave compartments of the greenhouse were regulated by an air-conditioning system (Kairos, Anjou Automation, Mortagne sur Sèvre, France). Temperatures were recorded at the pot level using a sensor (Aranet4, SAF Tehnika JSC, Riga, Latvia). Plants exposed to heatwaves were watered to the same extent (frequency and amount) as control plants. Additionally, we recorded the variation in pot mass before and after each heatwave event to evaluate any heat-induced water deficit. For each thermo-protocol, we had eight replicates per monoculture, twelve replicates of grass-legume mixture and sixteen replicates of four-species-mixture, making a total of 240 pots.

### Nutrient solutions

Throughout the experiment, plants were kept well-watered using a quarter-strength Hoagland solution consisting of: 1.25 mM Ca(NO□)□·4H□O, 1.25 mM KNO□, 0.25 mM KH□PO□, 0.5 mM MgSO□, 0.2 mM EDTA (NaFe·3H□O), 0.014 mM H□BO□, 0.005 mM MnSO□·H□O, 0.003 mM ZnSO□·7H□O, 0.0007 mM CuSO□·5H□O, 0.0007 mM (NH□)□Mo□O□□, and 0.0001 mM CoCl□. One half of the plants received this standard nutrient solution, while the other half was supplied with the same mineral concentrations except for MgSO□ which was 0.1 mM instead of 0.5 mM. Nutrient solutions were delivered via an automatic closed circulation system. The solution was replenished or replaced every three days, and each pot received a total of 300 mL per day, divided into five doses.

### Response variables

At the final harvest (July 1^st^), aboveground mass and root mass were assessed for each pot. The aboveground mass was split into two compartments i.e. above 5 cm (fodder harvest) and the rest. Biomasses were dried at 50°C to reach a constant mass and weighed in a precision scale (3200 g/0.01 g, MS3002S/01, Mettler Toledo, Columbus-OH, USA).

A subsample of the aboveground (above 5 cm) mass was finely ground using a ball mill (Retsch MM200, Restch GmbH, Haan, Germany) and the resulting powder was used for biochemical analyses. For each species grown in monoculture, we measured the percentage of Sulfur per mg DM. Sulfur content and δ13C were determined by a CNSOH element analyzer (EA Isolink, Thermofisher) linked to a continuous flow isotope mass spectrometer (Delta V advantage (Thermofisher). This elemental analysis was performed by Plateau technique d’isotopie de Normandie (PLATIN’). To measure leaf Sulphate content, 20□mg of ground leaf dry matter was used in a four-step extraction process. Leaf material was mixed with 0.5 ml of 50% ethanol and then incubated at 45°C for 1 h. After centrifuging the mixture for 10 min at 10 000g (4°C), the supernatant was collected and the pellet was re-extracted following the same procedure. The last two extractions were performed on the pellet with 0.5 ml distilled water at 95°C for 1 h (for each extraction). The final 2 ml of supernatant was concentrated for 20□hours at room temperature (Concentrator plus, Vacufuge® plus, Eppendorf), then suspended in 1 ml of ultrapure water and filtered (0.5 μm). The sulphate concentration was determined by High Liquid Performance Chromatography (HPLC, ICS-6000, Thermofisher). The eluent solution consisted of 4.05 mM Na2CO3 and 1.26 mM Na2HCO3, and was pumped isocratically at 1ml/min over an IonPac AG22 guard column 5 cm, 4 mm (Thermofisher 064139) and AS22 separation column 25 cm, 4 mm (Thermofisher 064141). To measure glutathione content, samples were grinded in liquid nitrogen. Extractions of glutathione were performed on 200 mg of samples with 500 µL of 5 % 5-sulfo-salicylic acid dihydrate (86192, Fluka) in an Eppendorf, a metallic ball was added in the tube and put in a grinder for 2 min, frequency 30/s (Retsch MM400). The supernatant was collected after centrifuging the mixture at 16000 g for 10 min (4°C). The total glutathione dosage was performed with the Glutathion Colorimetric Detection Kit (EIAGSHC, Thermofisher), in microplate, as described in the product information sheet. Samples were diluted to 1:5 with the Assay buffer provided in the kit and a second dilution was performed at 1:4 with the sample diluent described in the product sheet. The absorbances was red with a microplate reader (Tecan, Infinite M200 Pro) at 405 nm.

Heat-sensitive parameters were measured on the last day of the severe heatwave between 10 and 12 am. Leaf temperature and the maximum quantum efficiency of Photosystem II were measured on healthy leaves in the upper third of the canopy. Leaf temperature was measured at one point on the leaf with a thermal imaging camera (WB-80, Voltcraft, Hirschau, Germany). We measured one leaf for each species per pot for both the pots subjected to the mild and severe heatwaves and the control pots, in both Sulfur nutrition levels. The maximum quantum efficiency of Photosystem II was measured at one point on the leaf using a leafclip with an excitation fluorimeter (PocketPea, Hansatech Instruments, Norfolk, United Kingdom). The measured points were left 15 min in the dark before excitation with 2800 µmol m^-2^s^-1^ intensity. We used the Fv/Fm presented as a ratio of variable fluorescence (Fv) over the maximum fluorescence value (Fm) and calculated as (Fm - Fo)/Fv.

### Data analyses

To assess the effect of the thermoprotocol treatments, Sulfur nutrition and type of culture in the studied variables, we carried out ANOVAs using linear models with thermoprotocol treatments, Sulfur nutrition and type of culture as categorical fixed factor. We run linear models using glmmTMB function in glmmTMB package (Brooks *et al*., 2024). Heteroscedasticity and the normality of residuals were verified using Dharma package (Hartig and Lohse, 2022). We selected another residual’s family instead of Gaussian’s family when residuals were not normally distributed according to the type of variable.

## Results

### Shoot and root mass production

Shoot mass on studied grassland species was determined by the type of culture, Sulfur nutrition, and thermoprotocols. Significant effects were observed for the three factors alone, and the interactions thermoprotocol x Sulfur and thermoprotocol x Sulfur x culture (Table 1). S nutrition interacted with thermoprotocols in contrasting ways. Under S limitation, thermoprotocols of control, mild heatwaves and recurrent heatwaves had similar shoot mass, and thermoprotocol of severe heatwaves had the lowest value. While, under S standard level, the lowest was the thermoprotocol of recurrent heatwaves, when the other thermoprotocols had similar values. Overall, plants growing in S limitation (less those of severe heatwaves) had the highest shoot mass in our experiment jointly with plants exposed to mild heatwaves (S standard). Whereas, the least productive in our experiment were plants exposed to recurrent heatwaves (S standard) and plants exposed to a severe heatwave (S limitation) (Figure 1, Table S1). Regarding the type of culture, the most productive ones, regardless of Sulfur nutrition and thermoprotocols, were monoculture of *L. perenne* and grass-legume mixture, then the four-species mixture, monocultures of *P. lanceolata* and *F. rubra*, and the least productive was monoculture of *L. corniculatus*.

**Table 1.**
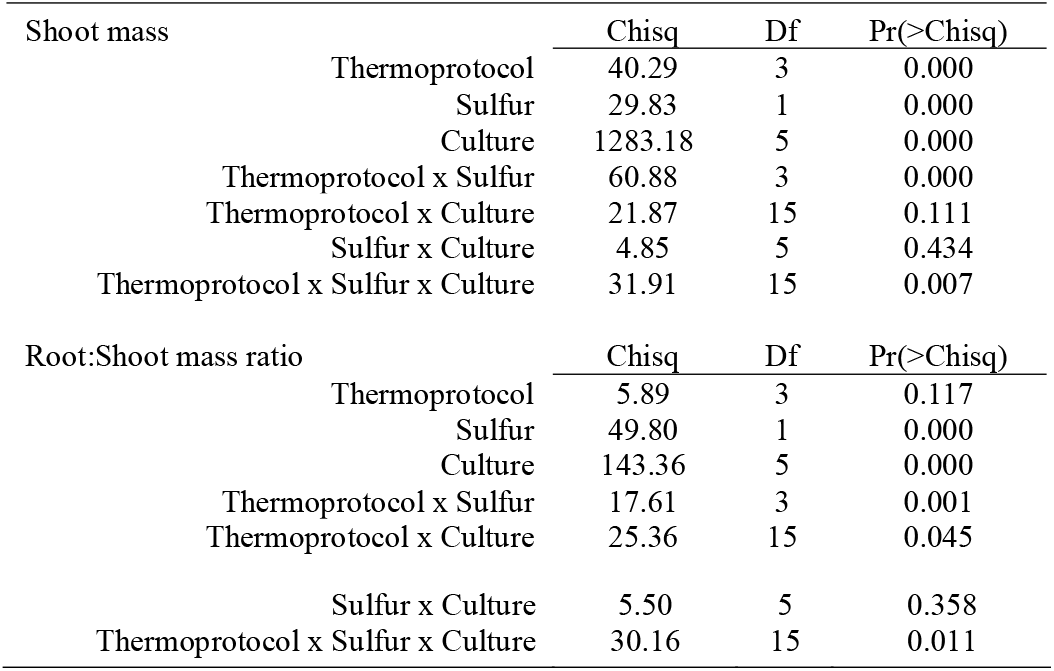
ANOVA summary of the effect of thermoprotocol, Sulfur nutrition and type of culture on shoot mass. Residual’s family for shoot mass was Gaussian, and for root:shoot mass ratio was Gamma.

| Shoot mass |  | Chisq | Df | Pr(>Chisq) |
| --- | --- | --- | --- | --- |
|  | Thermoprotocol | 40.29 | 3 | 0.000 |
|  | Sulfur | 29.83 | 1 | 0.000 |
|  | Culture | 1283.18 | 5 | 0.000 |
|  | Thermoprotocol x Sulfur | 60.88 | 3 | 0.000 |
|  | Thermoprotocol x Culture | 21.87 | 15 | 0.111 |
|  | Sulfur x Culture | 4.85 | 5 | 0.434 |
|  | Thermoprotocol x Sulfur x Culture | 31.91 | 15 | 0.007 |
| Root:Shoot mass ratio |  | Chisq | Df | Pr(>Chisq) |
|  | Thermoprotocol | 5.89 | 3 | 0.117 |
|  | Sulfur | 49.80 | 1 | 0.000 |
|  | Culture | 143.36 | 5 | 0.000 |
|  | Thermoprotocol x Sulfur | 17.61 | 3 | 0.001 |
|  | Thermoprotocol x Culture | 25.36 | 15 | 0.045 |

|  |  |  |  |
| --- | --- | --- | --- |
| Sulfur x Culture | 5.50 | 5 | 0.358 |
| Thermoprotocol x Sulfur x Culture | 30.16 | 15 | 0.011 |

**Figure 1.**
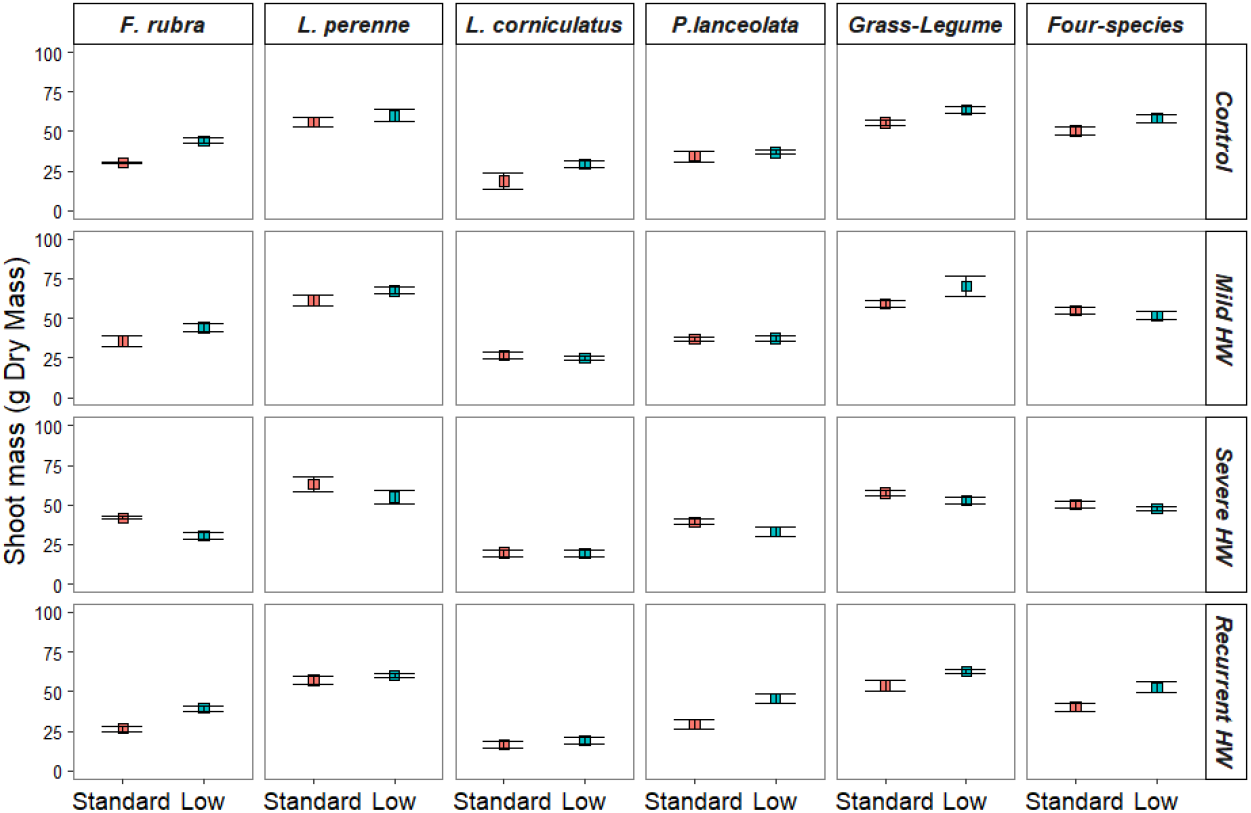
Shoot mass at the final harvest. Means and standard errors are shown. The vertical and horizontal panels distinguish respectively the types of cultures (monocultures and mixtures) and the thermoprotocols. For each combination (type and thermoprotocol), values are given for Standard and Low Sulfur nutrition levels.

Root:shoot mass ratio on studied grassland species and mixtures was determined by the type of culture, Sulfur nutrition, and thermoprotocols. Significant effects were observed for Sulfur nutrition and culture alone, and the interactions thermoprotocol x Sulfur, thermoprotocol x culture and thermoprotocol x Sulfur x culture (Table 1). The higher root:shoot ratios were obtained for plants grown in low Sulfur nutrition (Figure 2, Table S2), although significative differences between Sulfur nutrition levels were significant only under recurrent heatwaves. Regarding the type of culture, the highest root:shoot ratio was that of the monoculture of *L. perenne*, then monoculture of *F. rubra* and both mixtures, and the lowest root:shoot mass ratios were those of monocultures of *P. lanceolata* and *L. corniculatus*.

**Figure 2.**
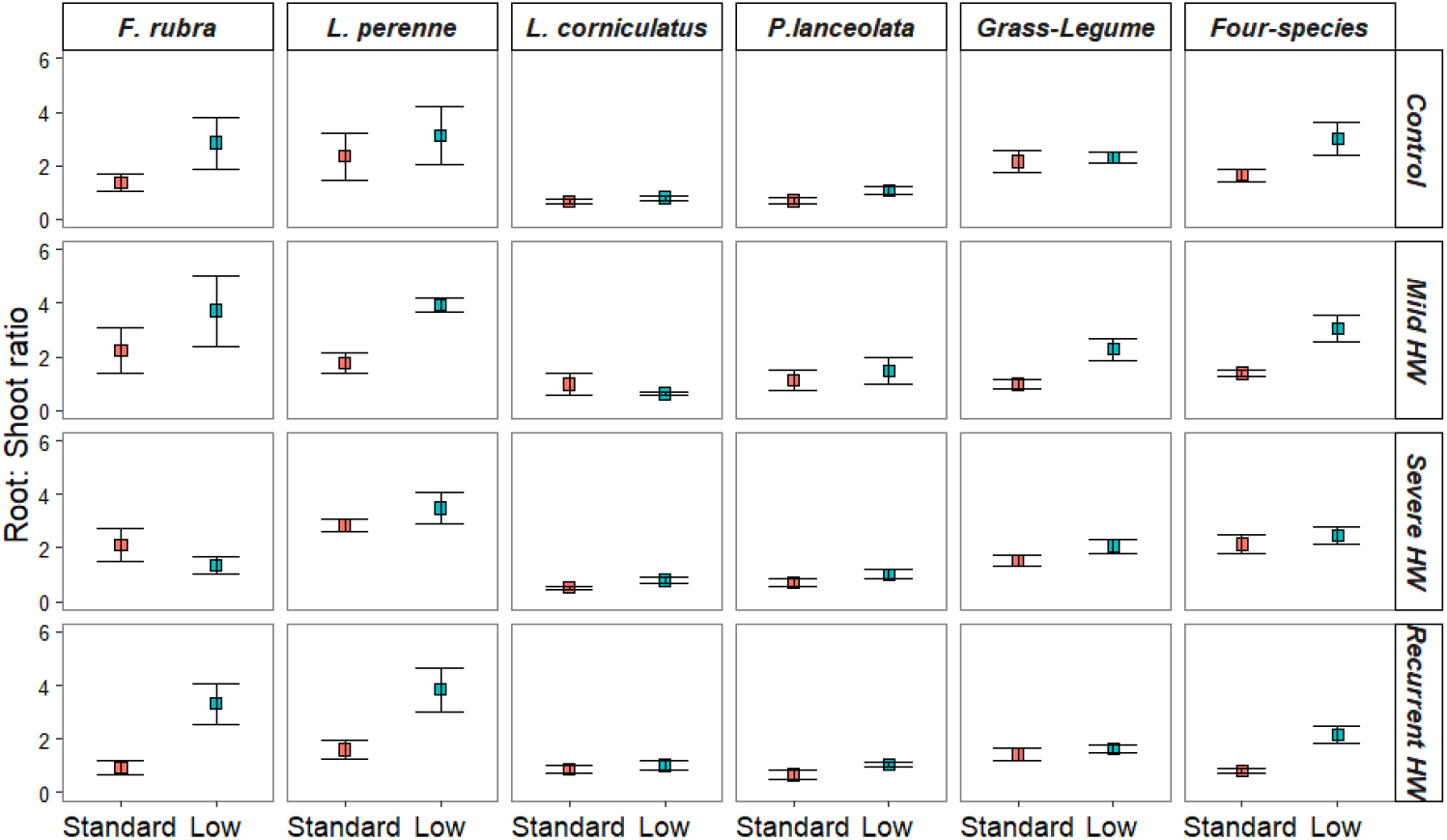
Root: Shoot mass ratio at the final harvest. Means and standard errors are shown. The vertical and horizontal panels distinguish respectively the types of cultures (monocultures and mixtures) and the thermoprotocols. For each combination (type and thermoprotocol), values are given for Standard and Low Sulfur nutrition levels.

### Sulfur, sulfate and glutathione concentration in shoot

In our experiment, sulfur and glutathione concentrations depended mainly on sulfur nutrition and the species. Thermoprotocol treatments alone did not have any effect on Sulfur, sulfate or glutathione concentrations (Table 2). For Sulfur and sulfate, significant effects were observed for Sulfur x species, and for the concentrations of the three compounds, Sulfur, sulfate and glutathione, the effect of the interactions thermoprotocol x species was also significant. The effect of the triple interaction was only significant for glutathione concentration. As expected, for all species, sulfur, sulfate (event not always significant) and glutathione concentrations were higher in plants growing in standard Sulfur nutrition than in low Sulfur nutrition (Figure 3). However, sulfur and sulfate concentrations varied among species according to Sulfur nutrition (Table S3, S4). In standard nutrition, *P. lanceolata* had the highest sulfur concentration, while *F. rubra* had the lowest sulfur and sulfate concentrations. In low nutrition, *L. corniculatus* had the highest sulfur concentration while *L. perenne* had the lowest, and all species had similar and very low concentrations of sulfate. For glutathione, regardless of sulfur nutrition and thermoprotocols, *L. corniculatus* had the highest concentrations (Table S5).

**Table 2.**
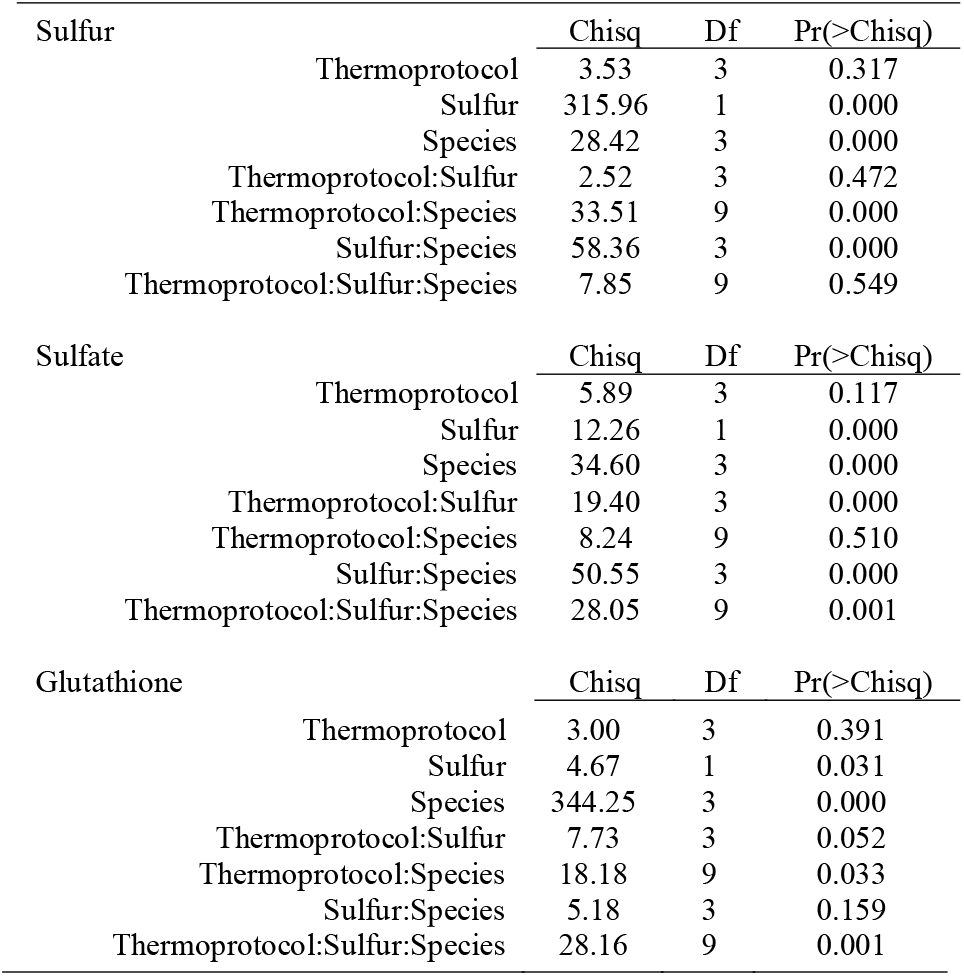
ANOVA summary of the effects of thermoprotocols, Sulfur nutrition and species on sulfur, sulfate and glutathione concentrations. Residual’s family for Sulfur concentration and glutathione was Gamma, and for sulfate ziGamma.

| Sulfur |  | Chisq | Df | Pr(>Chisq) |
| --- | --- | --- | --- | --- |
|  | Thermoprotocol | 3.53 | 3 | 0.317 |
|  | Sulfur | 315.96 | 1 | 0.000 |
|  | Species | 28.42 | 3 | 0.000 |
|  | Thermoprotocol:Sulfur | 2.52 | 3 | 0.472 |
|  | Thermoprotocol:Species | 33.51 | 9 | 0.000 |
|  | Sulfur:Species | 58.36 | 3 | 0.000 |
|  | Thermoprotocol:Sulfur:Species | 7.85 | 9 | 0.549 |
| Sulfate |  | Chisq | Df | Pr(>Chisq) |
|  | Thermoprotocol | 5.89 | 3 | 0.117 |
|  | Sulfur | 12.26 | 1 | 0.000 |
|  | Species | 34.60 | 3 | 0.000 |
|  | Thermoprotocol:Sulfur | 19.40 | 3 | 0.000 |
|  | Thermoprotocol:Species | 8.24 | 9 | 0.510 |
|  | Sulfur:Species | 50.55 | 3 | 0.000 |
|  | Thermoprotocol:Sulfur:Species | 28.05 | 9 | 0.001 |
| Glutathione |  | Chisq | Df | Pr(>Chisq) |

|  |  |  |  |
| --- | --- | --- | --- |
| Thermoprotocol | 3.00 | 3 | 0.391 |
| Sulfur | 4.67 | 1 | 0.031 |
| Species | 344.25 | 3 | 0.000 |
| Thermoprotocol:Sulfur | 7.73 | 3 | 0.052 |
| Thermoprotocol:Species | 18.18 | 9 | 0.033 |
| Sulfur:Species | 5.18 | 3 | 0.159 |
| Thermoprotocol:Sulfur:Species | 28.16 | 9 | 0.001 |

**Figure 3.**
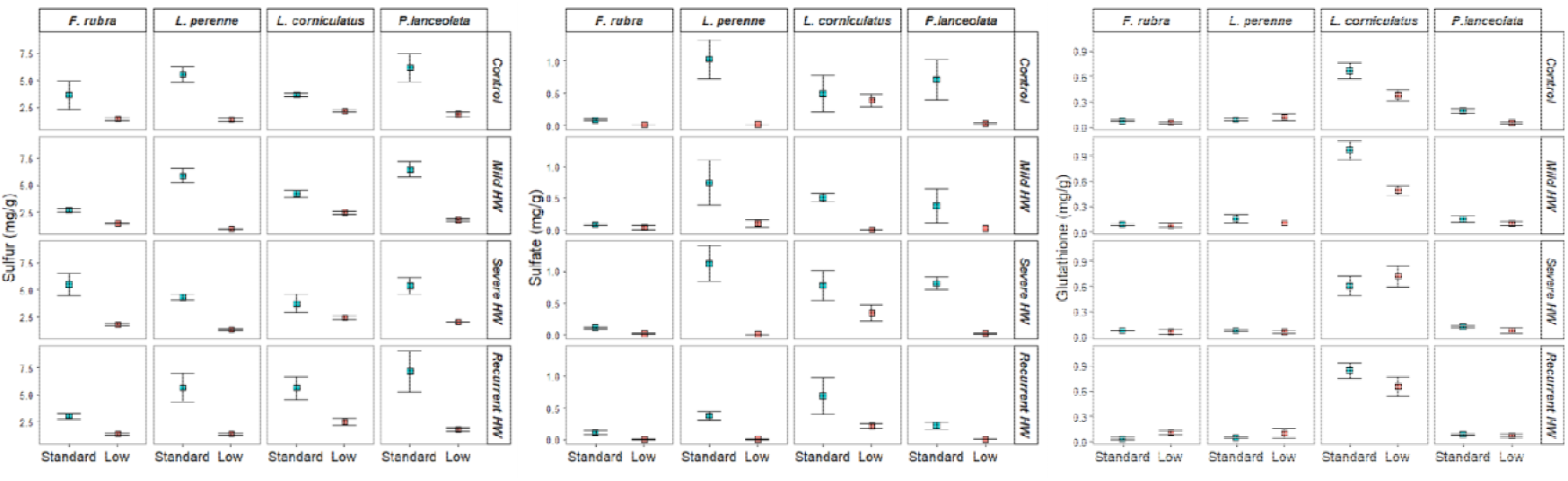
Sulfur, sulfate and glutathione concentrations at the final harvest. Means and standard errors are shown. The vertical and horizontal panels distinguish respectively the species monocultures and the thermoprotocols. For each combination (type and thermoprotocol), values are given for Standard and Low Sulfur nutrition levels.

### Physiological measurements sensitive to heatwaves

Values of δ^13^C, leaf surface temperature, and maximum efficiency of PSII varied according to the thermoprotocol treatments, Sulfur level and species, although effects on these variables were not all significant (Table 3). Significant species effects were observed on the three variables. Significant Sulfur effects were observed on the leaf surface temperature and there was a significant thermoprotocol x Sulfur interaction on δ^13^C and leaf surface temperature, and Sulfur x species interaction on δ^13^C (Table 3). Significant thermoprotocol effects alone were observed on the leaf surface temperature and the maximum efficiency of PSII. For δ^13^C, under severe heatwave alone, δ^13^C was higher under standard Sulfur nutrition than under low Sulfur nutrition, meaning that water use efficiency is higher under standard Sulfur than low Sulfur nutrition. Under recurrent heatwaves it was the opposite pattern (Figure 4, Table S6). In the case of leaf surface temperature, while plants under severe heatwave had similar values regardless of Sulfur nutrition, plants under recurrent heatwaves had lower leaf temperatures when growing in low Sulfur nutrition, regardless of species (Figure 4, Table S7). Values of maximum efficiency of PSII were different between thermoprotocol treatments with plants under recurrent heatwaves having the highest values, regardless of species (Figure 4, Table S8).

**Table 3.** ANOVA summary of the effects of thermoprotocols, Sulfur nutrition and species on δ^13^C, foliar surface temperature and maximum efficiency of PSII. Residual’s family for all variables was gaussian.

|  |  |  |  |
| --- | --- | --- | --- |
| $\delta^{13}\text{C}$ | Chisq | Df | Pr(>Chisq) |
| Thermoprotocol | 1.91 | 1 | 0.167 |
| Sulfur | 0.09 | 1 | 0.764 |
| Species | 10.39 | 3 | 0.016 |
| Thermoprotocol:Sulfur | 18.94 | 1 | 0.000 |
| Thermoprotocol:Species | 6.28 | 3 | 0.099 |
| Sulfur:Species | 10.89 | 3 | 0.012 |
| Thermoprotocol:Sulfur:Species | 7.79 | 3 | 0.051 |
| leaf surface temperature | Chisq | Df | Pr(>Chisq) |
| Thermoprotocol | 9.30 | 1 | 0.002 |
| Sulfur | 33.82 | 1 | 0.000 |
| Species | 19.10 | 3 | 0.000 |
| Thermoprotocol:Sulfur | 40.77 | 1 | 0.000 |
| Thermoprotocol:Species | 3.14 | 3 | 0.371 |
| Sulfur:Species | 0.24 | 3 | 0.970 |
| Thermoprotocol:Sulfur:Species | 1.50 | 3 | 0.682 |
| Maximum efficiency of PSII | Chisq | Df | Pr(>Chisq) |
| Thermoprotocol | 25.50 | 1 | 0.000 |
| Sulfur | 0.78 | 1 | 0.376 |
| Species | 62.26 | 3 | 0.000 |
| Thermoprotocol:Sulfur | 0.73 | 1 | 0.391 |
| Thermoprotocol:Species | 4.27 | 3 | 0.234 |
| Sulfur:Species | 0.18 | 3 | 0.980 |
| Thermoprotocol:Sulfur:Species | 0.96 | 3 | 0.812 |

**Figure 4.**
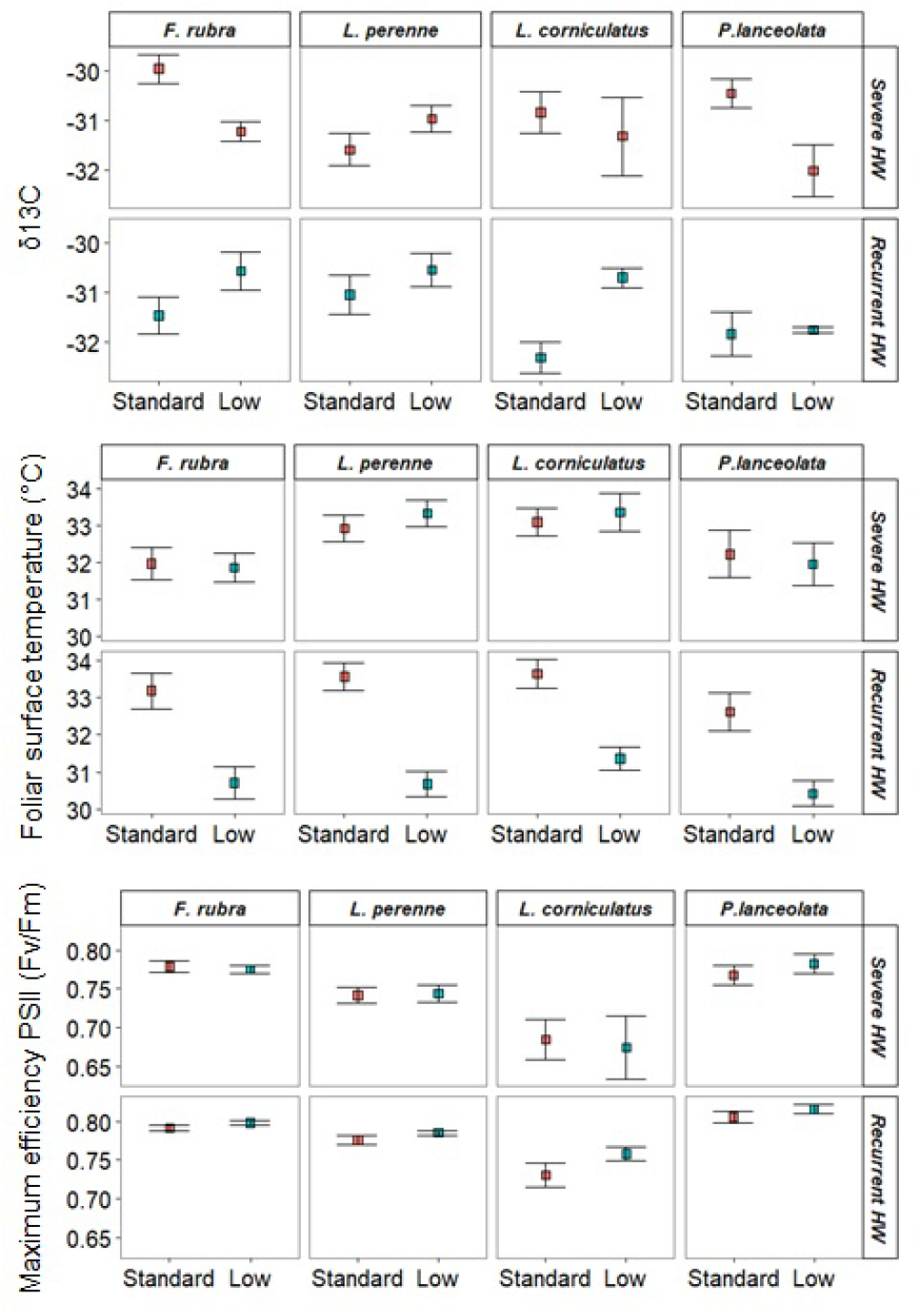
δ13C, foliar surface temperature, maximum efficiency PSII at the final harvest. Means and standard errors are shown. The vertical and horizontal panels distinguish respectively the species and the thermoprotocols. For each combination (type and thermoprotocol), values are given for Standard and Low Sulfur nutrition levels.

## Discussion

While we expected a pattern of reduced shoot mass and increased root:shoot mass ratio with Sulfur limitation, our results eventually show that the effects on plant growth varied according to the thermoprotocols. Although S limitation restricted S metabolism, as observed by lower concentration of S-compounds, this effect seemed to have no consequences on plant responses to heatwaves. On the contrary, S limitation induced indirect positive effects on plant responses to heatwaves, but only in the case of recurrent heatwaves, and not in the case of a single severe heatwave. Unexpectedly, these effects were overall similar for all cultures, both monocultures and mixtures.

### Plant growth under S limitation and heatwave exposure(s)

S limitation on grassland species have previously been shown to induce a reduction in shoot mass production (Tallec *et al*., 2009; Varin *et al*., 2009*a,b*; Cliquet and Lemauviel-Lavenant, 2019) and an increase in investment in root production, which results in an increase in the root:shoot mass ratio (Gilbert and Robson, 1984). However, we did not observe these effects completely, neither under control conditions nor under any thermoprotocol of heatwaves. We observed significant interactions between S levels and thermoprotocols on both shoot mass and root:shoot ratio. Overall, plants growing under low S nutrition showed higher shoot mass production than those growing with a standard level of S, but not when exposed to an isolated severe heatwave. Only under the recurrent heatwave thermoprotocol there was a clear investment in root production under S limitation. This result is interesting because the effect of heatwaves is known to depend on the intensity and frequency of the events (Cera *et al*., 2025) which determines the duration of the recovery period and so the ability of the plants to resist (Dreesen *et al*., 2014). In our case, these contrasting results between thermoprotocols may be due to the experimental design of S nutrition levels. We selected S limitation at 0.1 mM instead of S deficiency (at ∼0 mM) to be closer to realistic current soil availability in European grasslands (Scherer, 2001). Based on these results, we inferred that a mild heatwaves did not reverse the promotion of root production induced by S limitation, and even they could stimulated. Previous studies performed on herbaceous species have shown an increase in root production under heatwaves, which could be explained as a response to increased plant water demand associated with increased transpiration (Mainali *et al*., 2014) or better nutrient availability (Dreesen *et al*., 2012). In our experimental conditions, we observed this effect during the subsequent severe heatwave, which buffered shoot production due to the previous increase in root production. This ensured water availability even if we chose to avoid water stress and all plants were well watered with the same amount of water at the same frequency. This mechanism has already been evidenced for wheat under drought stress, with an increase in the root:shoot ratio improving plant water uptake and mitigating yield losses (Bacher *et al*., 2022). The improved response to a severe heatwave after having been under mild heatwaves can reflect acclimation. The change in the root:shoot ratio as an answer to the cumulative effects of heatwaves does indeed correspond to phenotypic plasticity (Gessler *et al*., 2020).

### Physiological regulations of heatwave exposure(s) effects

The indirect -through root growth promotion-positive effect of S limitation on shoot production under the thermoprotocol of recurrent heatwaves was demonstrated by the measured leaf surface temperatures during the severe heatwave and δ^13^C. Although plants rely on leaf cooling to reduce leaf temperatures during heatwaves (Urban *et al*., 2017), heatwaves can ultimately produce water stress if the heat persists (Körner, 2013). Interpreting leaf temperatures and δ^13^C values during heatwaves can provide a better understanding of plant responses to heatwaves. In our case, maximum PSII efficiency, a parameter frequently used in studies to interpret the effects of heatwaves (Geange *et al*., 2021), was not informative as the simulated maximum temperatures were below the plant’s thermotolerance threshold (Cera *et al*., 2025, Preprint). Plants growing under S limitation had lower leaf temperatures and higher WUE (i.e. fewer negative δ13C values) than those with standard S nutrition under the recurrent heatwaves thermoprotocol. However, under the severe isolated heatwave thermoprotocol, plants exposed to S limitation had similar leaf temperatures and lower WUE than those exposed to standard S nutrition. These results are consistent with those observed in shoot production and root:shoot mass ratio. S limitation only buffered plant responses to severe heatwaves under the thermoprotocol of recurrent heatwaves and not under the thermoprotocol of a severe isolated heatwave. Our interpretation is that plants that invested more in root production during previous heat episodes – owing to S limiting conditions-had better water regulation, resulting in higher WUE and lower foliar temperature during the severe heatwave.

### Direct effects of S nutrition on heatwave exposure(s) effects

The direct role of S and S-containing metabolites on plant stress management as previously described for other abiotic factors (Hawkesford *et al*., 2012) is less clear in our study. In our study, where we measured the concentrations of S, sulfate and glutathione, we observed that S limitation resulted in lower concentrations of S, sulfate and glutathione in leaves, but did not interact with the designed thermoprotocols. Leaf sulfate concentrations are tightly related to S availability, since plants remobilize foliar sulfate in the case of S limitation (Abdallah *et al*., 2010). In our case, leaf sulfate concentrations in low S nutrition were much lower than those observed under standard S nutrition and similar to those of low S nutrition in similar studies (Cliquet and Lemauviel-Lavenant, 2019; Kanté *et al*., 2022), indicating a clear S limitation. Regarding glutathione, which contribution to face a wide range of abiotic stresses has been largely documented es (Kopriva and Rennenberg, 2004), our experimental conditions did not highlight any significant contribution to the responses to heatwaves. It is important to raise that most studies that describe the role of S metabolism in response to abiotic stresses are short-term experiments (North and Kopriva, 2007; Gallardo *et al*., 2014), whereas our lasted 9 months. Therefore, integrating both (short and long terms) temporal approaches could allow the role of S-compounds in plant responses to heatwaves to be better analyzed.

## Conclusion

Our results have implications for interpreting grassland functioning under S limitation and heatwaves. Firstly, S limitation did not jeopardize plant responses to heatwaves, but rather improved them in certain condition. We did not observe any limitation to the positive effect of diversity in coping with heatwaves due to S limitation. Hence, interspecific competition for S did not appear higher than intraspecific competition. The promotion of root production stimulated by S limitation under mild heatwaves appears as an important indirect priming mechanism. Therefore, these results highlight the importance of taking a mid-term experiment to understand the effects of nutrient availability.

